# Hydroxide Once or Twice? A combined Neutron Crystallographic and Quantum Chemical Study of the Hydride Shift in D-Xylose Isomerase

**DOI:** 10.1101/030601

**Authors:** Matt Challacombe, Nicolas Bock, Paul Langan, Andrey Kovalevsky

## Abstract

The hydride shift mechanism of D-Xylose Isomerase converts D-glucose to D-fructose. In this work, we compute features of a “hydroxide once” mechanism for the hydride shift with quantum chemical calculations based on the 3KCO (linear) and 3KCL (cyclic) X-ray/neutron structures. The rigid boundary conditions of the active site “shoe-box”, together with ionization states and proton orientations, enables large scale electronic structure calculations of the entire active site with greatly reduced configuration sampling. In the reported hydroxide once mechanism, magnesium in the 2A ligation shifts to position 2B, ionizing the O2 proton of D-glucose, which is accepted by ASP-287. In this step a novel stabilization is discovered; the K183/D255 proton toggle, providing a ~10 kcal/mol stabilization through inductive polarization over 5Å. Then, hydride shifts from glucose-O2 to glucose-O1 (the interconversion) generating hydroxide (once) from the catalytic water. This step is consistent with the observation of hyroxide in structure 3CWH, which we identify as a branch point. From this branch point, we find several routes to the solvent-free regeneration of catalytic water that is strongly exothermic (by ~20 kcal/mol), yielding one additional hydrogen bond more than the starting structure. This non-Michaelis behavior, strongly below the starting cyclic and linear total energy, explains the observed accumulation of hydroxide intermediate – we postulate that forming permissive ionization states, required for cyclization, may be the rate limiting step. In all, we find eight items of experimental correspondence supporting features of the putative hydroxide once mechanism.

## 1. Introduction

Xylose Isomerase (XI) is responsible for aldose to ketose isomerization of hexose and pentose sugars via a hydride shift mechanism [1,2], and is one of the slowest enzymes known, remaining 5 orders of magnitude slower than triosephosphate isomerase that employs the ene-diol mechanism to catalyze a similar aldose to ketose conversion of triose sugars [3,4]. Despite its structural elucidation 25 years ago [5], its current industrial relevance [6,7], and the potential impact on the production of lignocellulosic ethanol [8–11], efforts to improve the activity of XI have yielded results primarily through enhancement of binding and stability [12–15], rather than through acceleration of the catalytic rate *k*_cat_. Currently, challenges to improving the catalytic rate are poorly understood, due to complexity of the mechanism, which involves a sugar ring opening step, together with an isomerization (interconversion) step that may involve non-Michaelis behavior [25,26]. In this work, we are concerned with the XI catalyzed interconversion of D-Glucose to D-Fructose as shown in Fig. 1.

**Figure 1.**
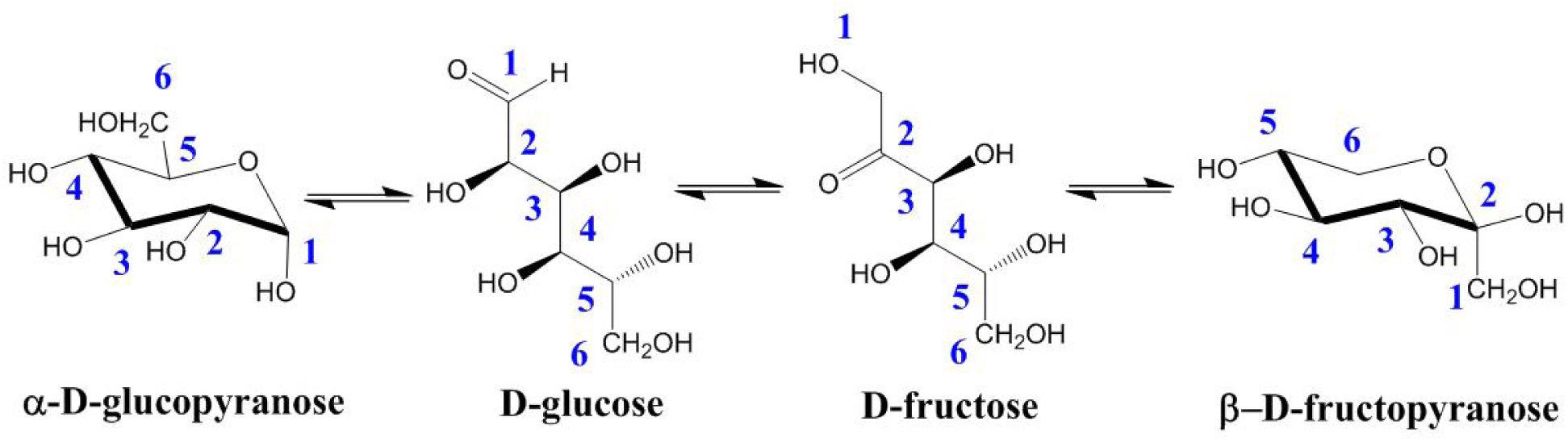
Conversion of D-Glucose to D-Fructose by Interconversion. The complete reaction catalyzed by XI is believed to involve three major steps: (1) ring opening, (2) isomerization, and (3) ring closure.

The isomerization step takes place in a bridged bimetallic active site that is enclosed in a hydrophobic “shoe-box” [14,16,17] enclosure of aromatics, including three tryptophane (W16+20+137), two histadine (H54+220) and two phenylalanine (F26+94) residues, which limit solvent exchange when the substrate (GLO401) is bound [18]. The active site consists of one metal position responsible for substrate binding, M1, and another with disorder (M2A/B [19] or M2a/b/c [3]) that is thought to initiate the hydride shift on movement, e.g. when the metal shifts from position M2A to position M2B.

High resolution X-ray studies involving Mn and xylitol find the Mn2· · · O_cat_ distance to be 2.4 Å at Mn2A and 1.8-2.0 Å at Mn2B (2a and 2b/c respectively in Ref. [[3]]). This result is interpreted as activation of the catalytic water 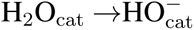, which remains in place as M2A shifts to M2B, with hydroxide later serving as the shift initiating base that deprotonates the O2 hydroxyl of the linear glucose [3,20–23]. In this interpretation, the hydroxide ion is generated twice, first in order to abstract the proton from O2 of glucose (perhaps involving an intermediate base such as D257 [3]), and again after the hydride shift, following the protonation of fructose O1.

However, recent neutron/X-ray structures of XI in complex with linear D-glucose (PDB ID 3KCO) [19] and with Ni^2+^ substitution at M1, M2A & M2B and shown in Fig. 2, find the catalytic water intact with distances M2B· · · O_cat_ = 2.0Å and M2A· · · O_cat_ = 2.7Å. Also, extended theoretical studies and crystallographic surveys by the Glusker lab demonstrate that a Mg· · · O bond length of ~2.0Å is consistent with octahedral coordination of Mg^2+^ [24]. Intriguingly, ionization of the catalytic water *is* found by neutron study in the 3CWH PDB structure [29] (*M* = Mg^2+^ and linear D-xylulose) with a 2.2Å Mg2A· · · O_cat_H^−^ bond length, ostensibly an intermediate trapped by kinetic or thermodynamic effects related to the extreme low reaction rate. These results suggest that the observed difference in M2B· · · O_cat_ and M2A· · · O_cat_ bond lengths may not correspond to a change in ionization of the catalytic water prior to isomerization, and that modes along the M2A → M2B axis are sloppy, with variations in observed distances corresponding perhaps to poorly understood ensemble differences in non-native metal substitutions.

**Figure 2.**
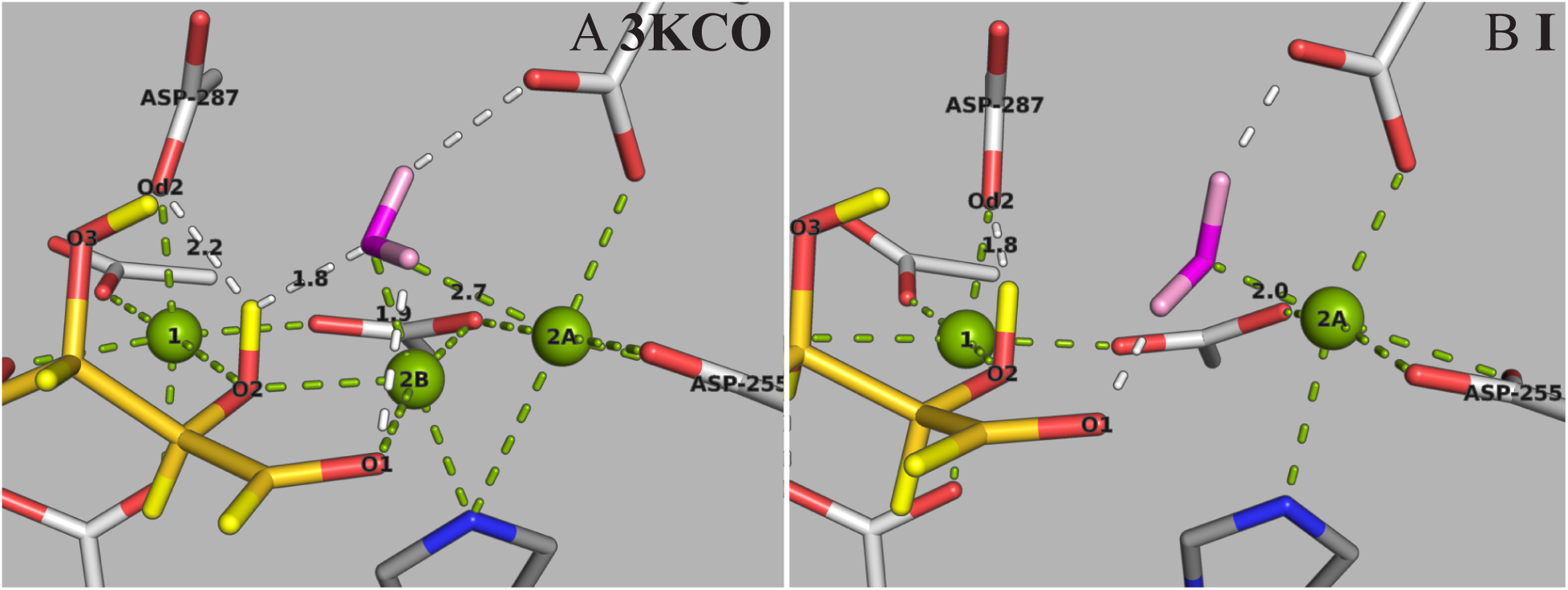
**A:** The Ni substituted 3KCO active site with the M1 (binding) and the M2A & M2B (disordered) metal positions in green, D-glucose in yellow and the catalytic water in magenta. **B:** The corresponding relaxed model structure, Model **I**, with bi-metallic Mg substitution at positions M1 and M2A.

Also, as shown in Fig. 2A, the “hydroxide twice” scheme would create the unshielded contact D287-O*δ*2^−^ … O2^−^-GLO401; its not clear how this contact can be compensated by coordinating ion-pairs, though it cannot be ruled out. Here we consider an alternative “hydroxide once” mechanism that does not involve an initiating base; instead the glucose-O2H proton is ionized directly by M2A-to-M2B movement as suggested by Collyer & Blow [25,26], with D287-O*δ*2^−^ accepting the initiating proton as argued by Fenn, Ringe & Petsko [3].

Previous theoretical work on XI has been based on X-ray data only, without knowledge of proton locations, with severely truncated quantum regions, and also based on many-parameter semi-empirical and QM/MM models [22,32,33]. In this work we consider additional constraints that allow to dramatically narrow the solution space of this problem, including (**a**) *ionization states and proton orientations* via neutron crystallography (**b**) averaged rigid crystallographic frameworks from multiple structures, (**c**) chemical and mutational observations from the literature, and (**d**) *high level quantum chemical treatment of the entire aromatic shoe-box* – a key boundary effect enabled by 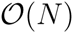 electronic structure theory. In this way, the present study is focused on representation of the primary (stiff) components in the mechanism that are constrained by the shoe-box boundary conditions.

## 2. Results

We present results for a “hydroxide once” mechanism of XI catalyzed D-glucose to D-fructose interconversion, with configurations obtaining from the 3KCO/3KCL basis, Section 4, via first order chemical transformations.

### 2.1. *Relaxation to Model* **I**

Starting with the boundary constrained crystallographic 3KCO model shown in Fig. 3A (described in Section 4), Mg is substituted for Ni in only the M1 and M2A positions (leaving the M2B site empty), corresponding to the chemically active bi-metallic occupation of XI. On relaxation to the corresponding enthalpy minimum, the rigorous octahedral coordination maintained by the 3KCO Ni ions distort, with Mg at the M1 position moving by 0.2Å and Mg at the M2A position moving by 0.3Å. Active site details are shown in Fig. 3A for the Ni substituted 3KCO model, and in Fig. 3B for the corresponding relaxed Model **I**.

**Figure 3.**
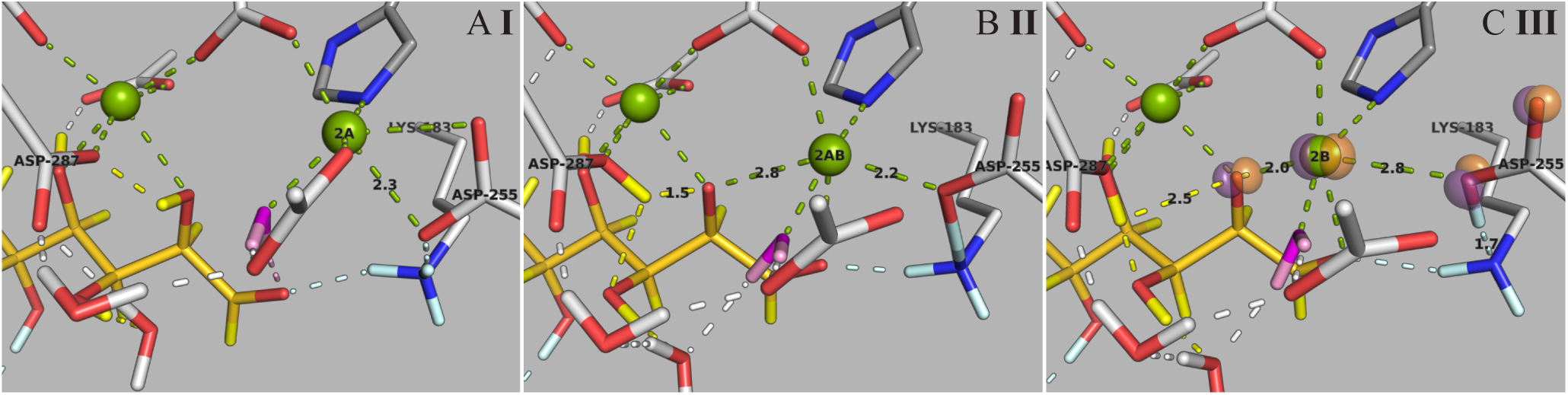
Mg shifts from position M2A to position M2B, ionizing the GLO401-O2 proton (yellow), which is accepted by D287-O*δ*2 (panels A & B). Concomitantly, a proton is transferred from 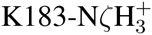 to D255-O*δ*2^−^ (light blue, panels B & C).

For Mg→ M2B, a barrier is confronted (Fig. 6), while with Mg about position M2A, the 3KCO contact GLO401-O2H· · · O_cat_ is lost with relaxation, and the catalytic water (magenta) shifts to establish a closer three-fold coordination with D257-O*δ*2, GLO401-O1 and Mg at position M2A (Fig. 3B), corresponding to a shortening of the M2A· · · O_cat_ distance from 2.7Å to 2.0Å. Also the H54-N*ζ*H^+^ proton is transferred to GLO401-O5. Details of Model **I** in context are shown in Vid. 8.

### 2.2. Ionizing the O2 Proton

Constrained optimization of Mg from M2A→M2B, described in Section 4, results in ionization of the GLO401-O2 proton to D287-O*δ*2, *and also proton transfer within the* 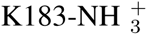 … O^−^-D255 *pair.* Optimization of this M2B intermediate yields Model **III**, shown in Fig. 3C. Transition state optimization connects basin **I** and **III** with TS Model **II**, shown in Fig. 3B, and with an enthalpic profile shown in Fig. 6. The reaction path for this transformation has the M1· · · M2 distance changing from 5.0 to 3.5 Å, with M2 moving in a plane defined by GLO401-O2, -C1 & -C2 and along the long axis of the shoe-box defined by the W16 & W20 rings.

Of particular interest is formation of the depolarized K183-H_2_N*ζ*· · · HO*δ*2-D255 ion pair, involving a 0.6 Å proton shift. Re-optimization of Model **III** with this proton constrained back to its native (Model **I**) bond length reveals an energetic cost of ~10 kcal/mol, as shown in Fig. 6. Also corresponding to this energy difference is an electrostatic stabilization potential, shown in Fig. 3C as ±0.2 au orange and purple isosurfaces. This stabilization potential is delocalized along the plane of M2 motion over ~ 5Å, including the centers GLO401-O2, M2 and D255-OO^−1^, as shown in Vid. 10. This stabilization corresponds to favorable quadrupolar polarizations about D255 and along the M2 … O2-GLO401 axis, corresponding to formation of the K183-H_2_N*ζ*· · · HO*δ*2-D255 ion pair.

### 2.3. Hydride Shift

The hydride shift involves transfer of a proton from GLO401-C2 to GLO401-C1 (D-Glucose → D-Fructose). In our model, the hydride shift is concerted with Mg returning to position M2A, reformation of the 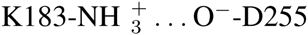 pair and the generation of 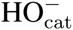 to arrive at Model **V**. This slightly hooked step includes the TS Model **IV** shown in Fig. 4 and in Vid. 11, with enthalpy profile given in Fig. 6.

**Figure 4.**
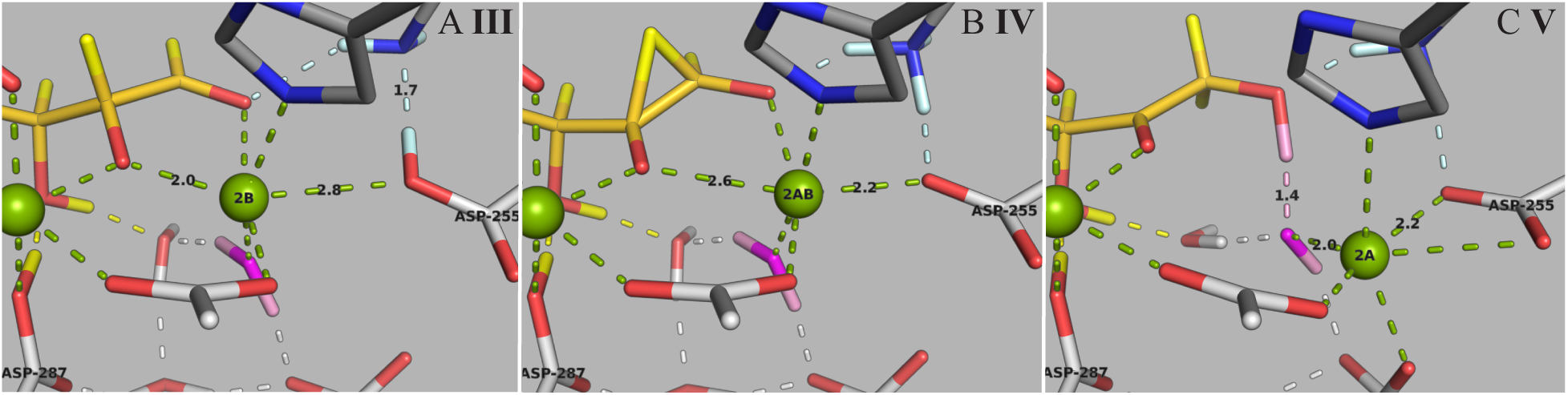
The hydride shift, concurrent with Mg moving towards position M2A and regeneration of the 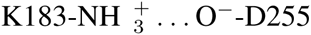 ion pair (panels A & B), followed by the formation of hydroxide by GLO401-O1 abstraction of proton from the catalytic water (pink, panels B & C).

### 2.4. Quenching Hydroxide

In the third step, the catalytic water is regenerated without solvent exchange through proton hopping mediated by the GLO401-O3 proton and water HOH1105 as shown in Fig. 5 and Vid. 12. This resolution however results in a substantial enthalpy drop of -20 kcal/mol, corresponding to formation of the contact HO_cat_H… O*δ*2-D287. Also, its worth noting the shallow plateau characterizing basin **V** 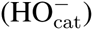, which branches also to regenerate the catalytic water via an alternative mechanism, involving proton hopping from 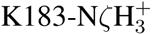 to the O1 of fructose as in Ref. [22], also with a substantial enthalpic drop comparable to the steps **V** → **VII**, *but resulting in a non-native ionization state that may (or may not) permit ring closure*.

**Figure 5.**
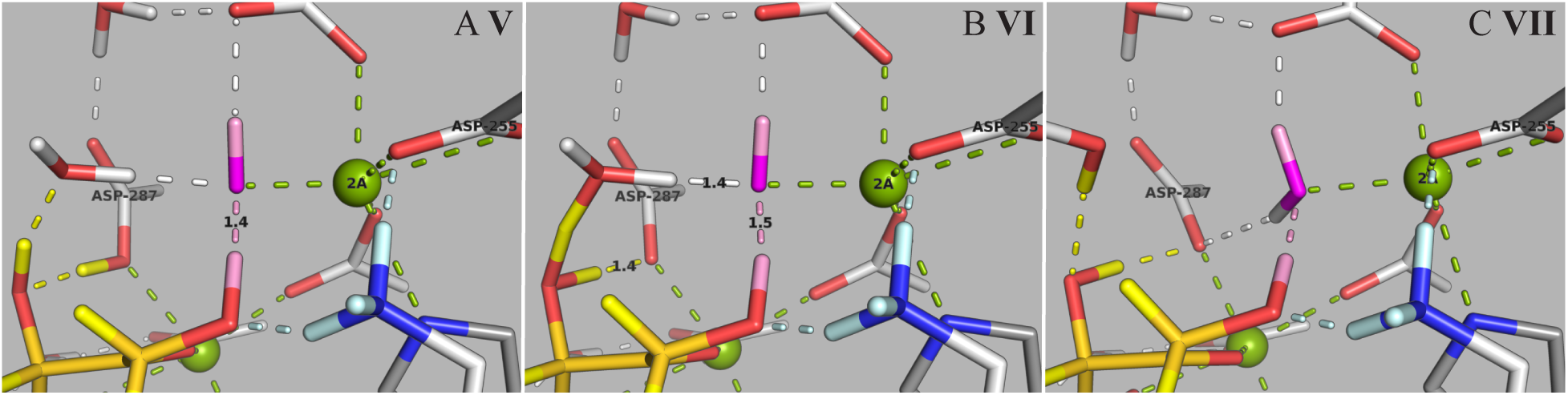
The ionized proton (D287*δ*O2-H, yellow) is reconciled with the catalytic hydroxide (pink) through proton hopping involving the proton GLO401O3-H (also yellow) and water HOH1105 (panels A & B), followed by exothermic regeneration of the catalytic water (pink). This energetic effect corresponds to the formation of an additional HO_cat_H… O*δ*2-D287 contact afforded by isomerization (panel D), resulting in 4 contacts for HO_cat_H relative to the starting 3, viz Fig. 2B.

## 3. Discussion

A primary approximation in this work is the substitution of three Ni atoms (M1, M2A and M2B) for two Mg atoms (M1 and M2A), viz Fig. 2, resulting in three primary contacts for the catalytic water, rather than the initial four, Fig. 2A vs. Fig. 2B. In Model I, the initial 2.7Å Mg· · · O_cat_ distance of the 3KCO basis is reduced to 2.0Å, consistent with other work characterization the Mg-O bond length [24,30].

Probing the M2A basin does not reveal additional, energetically accessible contacts for the catalytic water, and in the M2B case ionization of the GLO40-O2 proton follows, a result of Mg’s larger ionic radius in our interpretation (relative to Ni). Supporting acceptance of an ionized proton by D287, recent work demonstrates that the D287N mutant inactivates XI [15].

The K183/D255 toggle is a novel mechanism corresponding to inductive quadrupolar polarization, shown in Fig. 3C and in Vid. 10. This polarization is delocalized over ~5Å with a corresponding stabilization of ~10 kcal/mol. The K183/D255 toggle mechanism is supported by previous mutagenesis studies, including findings of reduced *k*_cat_ with G219F and G219N mutations that may sterically disrupt the ion pair [14], and inactivating mutations K183S, K183Q, K183R that either remove or dramatically reduce this charge transfer [16].

Our models of the hydride shift, **III** → **V** shown in Fig. 4 and Vid.11, are consistent with Allen *et al.*’s [28] structure of the putative transition state analogue D-threonohydroxamic acid (THA) complexed with XI; we also find a M1· · · M2 distance of 4.1Å at the transition state, viz Model **IV** of Fig. 4B. Also compelling is discovery of the hydroxide intermediate, Model **V**, following automatically from constrained optimization of hydride from GLO401-O2 to GLO401-O1 in the Model **III** basis; hydroxide is observed as a putative intermediate in the 3CWH structure [29] (*M* = Mg^2+^ and linear D-xylulose).

Shown in Fig. 6, our energetic results have three features consistent with observation: First, we find two, partially limiting enthalpic barriers of ~16 kcal/mol, in reasonable agreement with experiment (14 kcal/mol [1]). Second, the large enthalpic drop on isomerization is exothermic, consistent with non-Michaelis behavior [25,26]. Third, kinetic isotope effect measurements utilizing deuterium-labeled substrates or D2O suggest a partial rate limiting behavior between two steps: (**1**) a pH sensitive step (perhaps related to the K183/D255 pair) and (**2**) the hydride shift [2]. The partial limiting behavior of these two steps is consistent with the proposed mechanism as well as the two equi-enthalpic peaks corresponding to structures **II** and **IV**.

**Figure 6.**
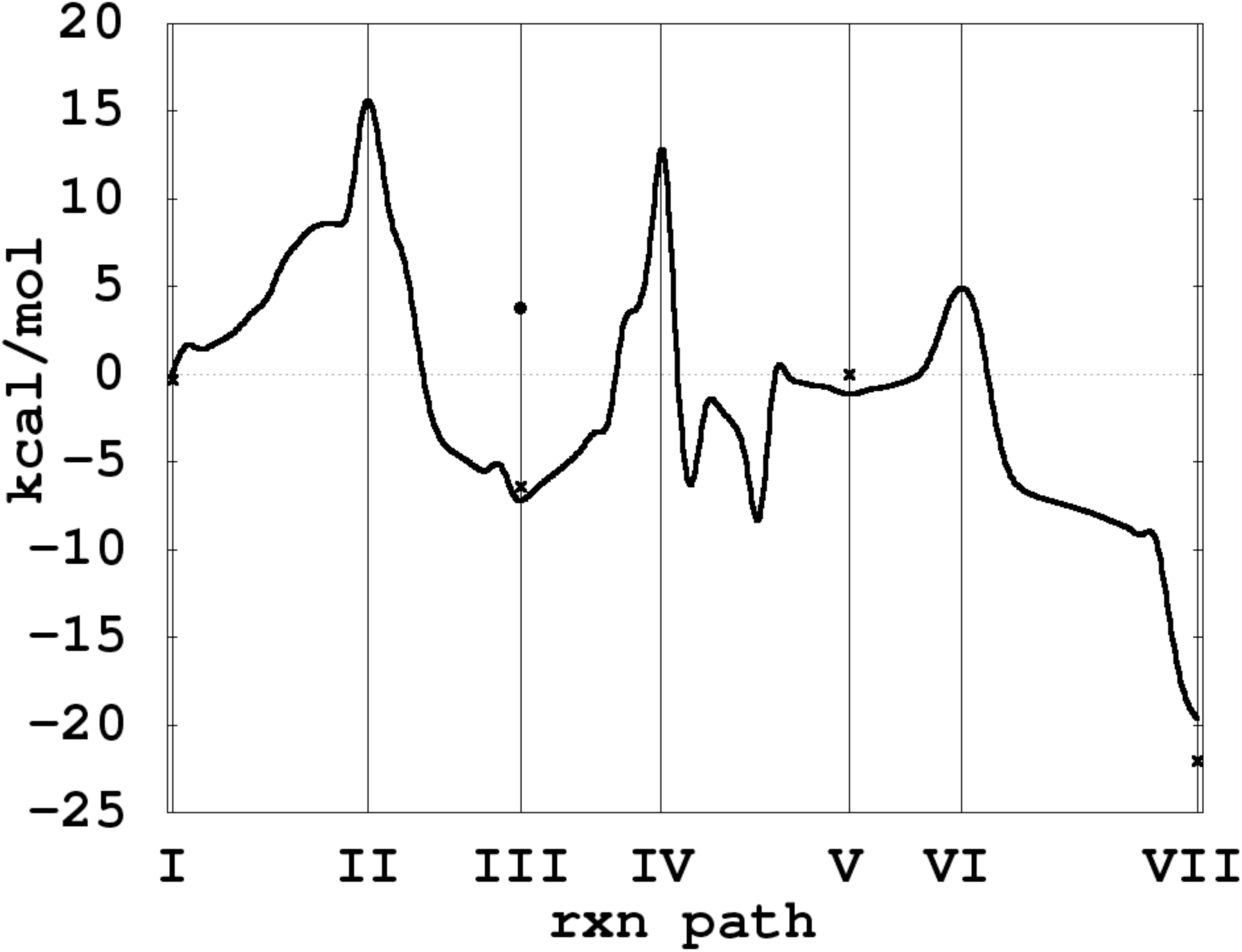
Enthalpic reaction profile for the putative “hydroxide once” mechanism, computed at the B3LYP/6-31G** level of theory, with larger B3LYP/6-311G** calculations at labeled minima marked by ×s. Also shown (•) is the energy difference for structure **III** with the K183-H_2_N*ζ*· · · H bond length constrained back to its original 0.98Å, corresponding to the stabilizing polarization of the electronic potential shown in Fig. 3C.

In this work, we have considered only first order changes associated with optimization into new basins along only one internal degree of freedom; Mg from M2A → M2B, and hydride from GLO401-O2 to GLO401-O1. These first order constraints lead to surprising concerted behaviors, like the K183/D255 toggle (**I** → **III**) and the formation of hydroxide (**III** → **V**). From **V** onward however, our model is more speculative, involving at least two alternative paths to regenerate the catalytic water, each with roughly similar energetics that are associated with quad-coordination of the catalytic water, and that do not require solvent exchange. The shallow basin **V** is a branch for these two (and possibly other) states, which is seemingly observable in the the 3CWH structure [29].

Unfortunately, we cannot argue too much further about limiting behavior in XI *ab initio;* although ionization of the GLO401-O2 proton and the hydride shift do not seem excessively large, it remains to work out the ring unfolding and folding, *e.g.* connecting the 3KCL and 3KCO basis. In preliminary work however, we find no large energy differences between cyclic and linear forms of D-Glucose, and in early parts of the ring opening involving proton transport, there are only low barriers. Also, the observed limiting steps, **1** & **2** above, are in reasonable agreement with the present results and alone cannot explain the observed extreme slow rate of XI.

**Table 1.** Items of correspondence between the “hydroxide once” model and experiment.

| Model | Observed | Refs. |
| --- | --- | --- |
| Mg⋯O<sub>cat</sub> distance is 2Å throughout | Mg⋯O ∼ 2Å | 1S5N[3], [24,30] |
| Hydroxyl ionization initiated by M2 and the hydride shift are both rate limiting steps. The K183/D255 toggle is pH sensitive and stabilizing by 10 kcal/mol | Two partially limiting steps: the hydride shift and another pH sensitive step | [2] |
| K183/D255 ion-toggle favorable by 10 kcal/mol as Mg moves from position M2A to position M2B. | Steric impedance of mutations G219F, G219N lower $k_{cat}$ while K183 S, K183Q & K183R eliminate proton transfer to D255, inactivating the enzyme | [16] [14] |
| D287-2δO<sup>-</sup>1 accepts ionized GLO401-O2 proton | D287N inactivates | [15] |
| 16 kcal/mol $\Delta H^\ddagger$ | 14 kcal/mol $\Delta G^\ddagger$ | [1] |
| Mg<sub>1</sub>⋯Mg<sub>2</sub> distance is 4.1Å in hydride shift transition state | Mg<sub>1</sub>⋯Mg<sub>2</sub> is 4.1Å with transition state analogue THA | 2GYI [28] |
| Non-Michaelis energetics | Non-Michaelis kinetics | [25,26,29] |
| Hydroxide intermediate | Hydroxide intermediate | 3CWH [29] |
| No solvent exchange | Shielded active site (bound substrate) | [18] |

### 3.1. Summary

In Table 1, we summarize the above eight items of correspondence, between the chemistry of XI and features of our computed reaction path, together with fourteen supporting references. Also, in Fig. 7 we present a summary diagram of the complete seven step reaction path of the putative “hydroxide once” mechanism developed here.

**Figure 7.**
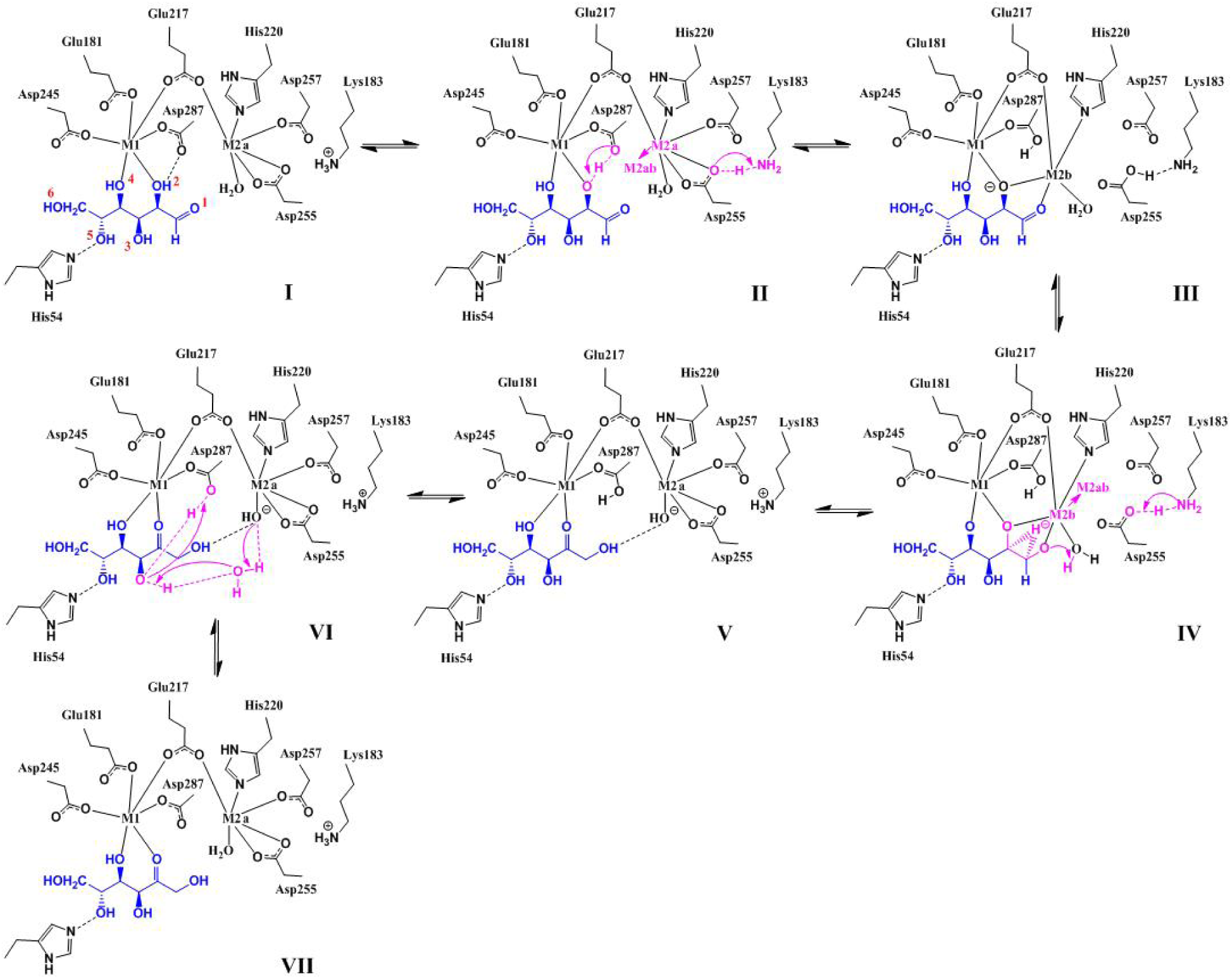
Diagrammatic representation of the proposed “hydroxide once” mechanism of D-glucose isomerization, corresponding to structures **I**-**VII** in Figs. 3-5, and the energy diagram in Fig. 6. Figure **V** is a branch point; steps **V**→**VII** are representative also of other near lying states that lead to solvent free regeneration of the catalytic water, involving for example 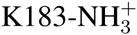 instead of D287-*δ*2OH.

## 4. Materials and Methods

### 4.1. *Model* **I** *in the 3KCO/3KCL Basis*

This work is based on the RMSD alignment and average of the rigid shoe-box structure bounding the solvent shielded active site of Xylose Isomerase (XI), corresponding to coordinates from both the 3KCO (linear sugar) and 3KCL (cyclic sugar) structures [19]. The shoe-box is strongly overlapping in these two configurations, suggesting a rigid boundary unperturbed by the cyclic → linear transformation. The shoe box is delimited by the aromatic rings of tryptophane (W16+20+137), histadine (H54+220) and phenylalanine (F26+94).

In our present model of XI, hard constraints are imposed on 101 heavy atoms at the truncated, methylated periphery, with an interior containing 205 mobile centers corresponding to the unaveraged 3KCO configuration, shown in Fig. 2A. On substitution of Mg at the M1 and M2A positions, the model is charge neutral, involving 306 total atoms including W16+20+137, H54+220, D245+255+257+287, K183+289, E181+217, Q256, T90+91, N92, F26+94, Mg391+392, GLO401 (D-glucose) & internal waters HOH1001+1095+1105+1106+1159+1190+1193+1163+1209. Our relaxation protocol, starting from this 3KCO/3KCL basis, involved fixing all interior heavy atoms and first relaxing protons, then relaxing waters and heavy atoms except those of the sugar, and finally full relaxation of all interior atoms. The resulting initial condition is shown in Fig. 2B and in Vid. 8, with the main features of this model described in Section 2.1.

### 4.2. Electronic Structure Calculations

All calculations were carried out using a serial (single CPU) version of the `freeon`^1^ suite of 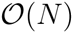 electronic structure codes [36]. In this version of `freeon`, fast methods rely on matrix sparsity but introduce a numerical uncertainty that depends on basis set ill-conditioning and on numerical truncation associated with small contributions to the total energy (enthalpy). In all calculations we used the `tight` threshold levels enabling the convergence of gradients to within 0.2 ev/Å, corresponding to a precision of 2 significant digits in reported energy differences at the RB3LYP/6-31G** level of theory [34,35]. For each labeled basin, we also checked these results with additional calculations at the higher RB3LYP/6-311G** level of approximation (× in Fig. 6).

All relaxations were carried out with the QUICCA optimizer [27], with bond constrained optimization enabling simple chemical transformations along principle internal coordinates. In arriving at steps **I** → **VII**, alternative but energetically less favorable states were also explored in this way, but not exhaustively so. Between tightly optimized basins **I**, **III**, **V** & **VII** transition state structures **II**, **IV** and **VI** were obtained through linear synchronous transit followed by iteration with the climbing image version of the Nudged Elastic Band (NEB) [37]. In the case of Model **III**, the K183-H_2_N*ζ*· · · H bond length was constrained back to its original 0.98Å using QUICCA, and the difference in electrostatic potential between constrained and unconstrained systems was computed (shown in Fig. 3C). In this way, the reaction profile shown in Fig. 6 was generated.

## 5. Conclusions

In this work we focused on the plain B3LYP/6-31G**/6-311G** quantum chemistry (bare enthalpy) of the hydride shift in Xylose Isomerase, with new configurational basins explored by only first order perturbations along sequential, atomic transformations (system optimization with one bond constraint), revealing novel behaviors along each transformation, including the K183/D255 toggle and the formation of hydroxide. This minimalist approach is possible due to the 3KCL and 3KCO configurational basis provided by the neutron/X-ray crystallography of Kovalevsky, Langan, Glusker and co-workers [19]. From this structural work, we know that the shoe-box is rigid under the perturbation of ring opening; we also have ion-pair charge states and also orientation of the internal water network, although experimental characterization of the catalytic water remains enigmatic; based on Models **I** and **III**, and hints from the difference FO-FC electron density maps in several ultra-high resolution X-ray structures [3,4,31], we conjecture that multiple states (A/B or a/b/c) exist for H_2_O_cat_.

From this configurational basis and using only modest (serial, single CPU) server resources, it was possible to correlate eight features of XI chemistry with properties of the *ab initio* reaction path, viz Table 1, with approximations only in (a) metal substitution, (b) model chemistry (bare B3LYP/6-31G**/6-311G**) and (c) shoe box boundary conditions. Without the boundary conditions provided by neutron crystallography, the size advantage obtained by 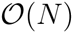 electronic structure would be rapidly diminished by the effects of dimensionality and associated demands on configurational sampling.

In addition to discovery of the K183/D255 toggle, a major finding in this work is the additional contact made by catalytic water on the solvent free resolution of hydroxide; four h-bonds in Model **VII** vs. three in Model **I**, together with the corresponding large enthalpy drop (*e.g*.~20 kcal/mol) below the putative starting point^2^. This non-Michaelis effect related to the build up of intermediate was observed already in 1990 by Collyer and Blow [25], as well as more recently by Kovalevsky *et al* as specifically the hydroxide intermediate (3CWH)[29]. From **V**, multiple exothermic paths to a solvent free quench may exist that involve unproductive ionization states in the active site, *i.e.* they are unfavorable to ring folding. We conjecture that the slowness of XI may be tied to quench states that do not favor ring folding, and that the enzyme may remain combinatorically and energetically trapped by these effects with transitions passing unproductively through **V**. Supporting this view, much faster enzymes that catalyze small sugar isomerization, such as triosephosphate isomerase, are unencumbered by ring opening; in XI, this ring opening is mediated by an extended internal (solvent shielded) network of waters and ion-pairs that may be susceptible to ionization states unfavorable to reforming the cycle.

**This work was supported by the U. S. Department of Energy under Contract No. DE-AC52-06NA25396 and LDRD-ER grant 20120256ER. References**

## 6. Supporting Information

**Figure 8.** Model **I**, the “shoe-box” active site of Xylose Isomerase. Model **I** is shown as thin and full sticks, in context of the XI protein backbone (ribbons). In later images and videos, only the full sticks are shown for clarity.

**Figure 9.** Mg shifts from position M2A to position M2B, ionizing the GLO401-O2 proton (yellow), which is accepted by D287-O*δ*2. Concomitantly, a proton is transferred from 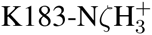 to D255-O*δ*2^−^ (light blue).

**Figure 10.** Details of the stabilizing polarization effect associated with the K183/D255 toggle, along the plane of ionization.

© 2016 by the authors; licensee MDPI, Basel, Switzerland. This article is an open access article distributed under the terms and conditions of the Creative Commons Attribution license (http://creativecommons.org/licenses/by/4.0/).

**Figure 11.** The hydride shift, concurrent with Mg moving towards position M2A and regeneration of the 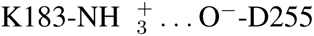 ion pair, followed by the formation of hydroxide by GLO401-O1 abstraction of proton from the catalytic water (magenta).

**Figure 12.** The ionized proton (D287*δ*O2-H, yellow) is reconciled with the catalytic hydroxide (pink) through proton hopping involving the GLO401-O3 proton (also yellow) and water HOH1105, followed by exothermic regeneration of the catalytic water (pink). This energetic effect cooresponds to the formation of an additional HO_cat_H… O*δ*2-D287 contact afforded by isomerization, resulting in 4 contacts for HO_cat_H relative to the starting 3, vis Fig. 2B.

## Footnotes

1 Version `beta:ad3a92337b33bef270e2288173fe3b804df07d54`

2 Corresponding to the “zero” in both the cyclic 3KCL and the linear 3KCO basis (unpublished).

